# Dnmt3b deficiency in adipocyte progenitor cells ameliorates obesity in female mice

**DOI:** 10.1101/2025.01.31.635994

**Authors:** Yifei Huang, Sean Yu, Qiang Cao, Jia Jing, Weiqing Tang, Bingzhong Xue, Hang Shi

## Abstract

Obesity arises from chronic energy imbalance where energy intake exceeds energy expenditure. Emerging evidence supports a key role of DNA methylation in the regulation of adipose tissue development and metabolism. We recently discovered a key role of DNA methylation, catalyzed by DNA methyltransferase 1 or 3a (Dnmt1 or 3a), in the regulation of adipocyte differentiation and metabolism. Here, we aimed to investigate the role of adipocyte progenitor cell Dnmt3b, an enzyme mediating *de novo* DNA methylation, in energy metabolism and obesity. We generated a genetic model with *Dnmt3b* knockout in adipocyte progenitor cells (PD3bKO) by crossing *Dnmt3b*-floxed mice with platelet-derived growth factor receptor alpha (Pdgfrα)-Cre mice. *Dnmt3b* gene deletion in adipocyte progenitors enhanced thermogenic gene expression in brown adipose tissue, increased overall energy expenditure, and mitigated high-fat diet (HFD)-induced obesity in female mice. PD3bKO mice also displayed a lower respiratory exchange ratio (RER), indicative of a metabolic shift favoring fat utilization as an energy source. Furthermore, female PD3bKO mice exhibited improved insulin sensitivity alongside their lean phenotype. In contrast, male PD3bKO mice showed no changes in body weight but demonstrated decreased insulin sensitivity, revealing a sexually dimorphic metabolic response to *Dnmt3b* deletion in adipocyte progenitor cells. These findings underscore the critical role of Dnmt3b in regulating energy homeostasis, body weight, and metabolic health, with significant implications for understanding sex-specific mechanisms of obesity and metabolism.

## Introduction

Obesity is closely associated with a panel of metabolic disorders such as type 2 diabetes, hypertension, dyslipidemia, and cardiovascular diseases (1). Obesity arises from a chronic energy influx due to energy intake exceeding expenditure (1). Thus, a better understanding of the mechanism governing energy metabolism may provide therapeutic strategies for the treatment of obesity and related metabolic disease.

Adipose tissue plays a central role in regulating energy homeostasis by balancing energy storage, mobilization, and dissipation. This dynamic process is mediated by three distinct types of adipocytes: white, brown, and beige. White adipose tissue (WAT) stores excess energy as triglycerides through hypertrophy, hyperplasia, or both, and releases fatty acids via lipolysis to fuel other organs during energy demand (2). In contrast, brown adipose tissue (BAT) dissipates energy through adaptive thermogenesis, employing both UCP1-dependent and -independent mechanisms (3–6). Beige adipocytes, sporadically dispersed in WAT depots, are primarily induced by β-adrenergic stimulation triggered by cold exposure or β-adrenergic receptor agonists (7). Sharing morphological and biochemical features with BAT, beige adipocytes also contribute to thermogenesis (7). Together, these adipocyte types perform distinct yet complementary roles in maintaining energy homeostasis.

Obesity, like many other complex diseases, arises from the interplay between genetic and environmental factors, such as diet. One mechanism by which environmental factors influence gene expression is through epigenomic reprogramming, a critical molecular link between obesity and environmental factors (8,9). DNA methylation, a common epigenetic mechanism, involves the covalent addition of a methyl group to cytosine, often at CpG sites. This modification typically occurs in gene promoters and 5’ regions, where CpG sites are enriched (10,11). Hypomethylation in promoter regions generally activates gene transcription, while hypermethylation silences genes by disrupting the binding of transcriptional activators or by cooperating with histone modifications to alter DNA accessibility (11,12). Three functional DNA methyltransferases (DNMTs), namely DNMT1, DNMT3A, and DNMT3B, catalyze DNA methylation in distinct contexts (11). DNMT1, which prefers hemimethylated DNA, primarily maintains DNA methylation patterns during replication. In contrast, DNMT3A and DNMT3B are responsible for *de novo* methylation, establishing new methylation patterns (11). However, emerging evidence suggests that DNMT1 may also contribute to *de novo* methylation under certain circumstances (13).

We recently identified DNA methylation as a key regulator of adipocyte development and metabolism. Specifically, we discovered that DNA methylation, catalyzed by DNMT1 and DNMT3A, has a biphasic role in 3T3-L1 cell differentiation: promoting adipogenesis during the early stages while inhibiting lipogenesis at later stages (14,15). Moreover, we found that DNMT1, DNMT3A and DNMT3B in brown fat are critical for regulating thermogenesis and diet-induced obesity in mice (16–18). In this study, we aimed to investigate the role of Dnmt3B in adipocyte progenitor cells and its impact on energy metabolism in adipose tissue. To achieve this, we generated a genetic model with *Dnmt3b* knockout in adipocyte progenitor cells (PD3bKO) by crossing *Dnmt3b*-floxed mice with platelet-derived growth factor receptor alpha (Pdgfrα)-Cre mice. We then characterized the metabolic phenotype of these mice under a high-fat diet (HFD).

## Materials and Methods

### Animals and diets

PD3bKO mice were generated by crossing *Dnmt3b*-floxed mice (Mutant Mouse Regional Resource Centers (MMRRC), stock # 029887) with Pdgfrα-Cre mice (Jackson Laboratory, Stock # 013148). The *Dnmt3b*-floxed mouse was created by inserting two loxP sites flanking exons 16-19 encoding the catalytic motif (19), and has been backcrossed to B6 background for more than five generations. All animal procedures in this study were approved by the Institutional Animal Care and Use Committee at Georgia State University (GSU). Mice were housed in a temperature- and humidity-controlled facility at GSU under a 12-hour light/dark cycle with free access to food and water. PD3bKO mice and their flox/flox (fl/fl) littermate controls were fed either a chow diet or a HFD (Research Diets D12492, 60% calories from fat) for up to 24 weeks.

### Metabolic analysis

During the HFD feeding study, body weight was measured weekly, and food intake was monitored in a single cage over seven consecutive days. Body composition, including fat and lean mass, was analyzed using a Minispec NMR body composition analyzer (Bruker BioSpin Corporation, Billerica, MA). Energy expenditure parameters, such as oxygen consumption, carbon dioxide production, and locomotor activity, were measured using the PhenoMaster metabolic cage system (TSE Systems, Chesterfield, MO). Blood glucose levels were measured with a OneTouch Ultra Glucose meter (LifeScan, Milpitas, CA), and glucose tolerance test (GTT) and insulin tolerance test (ITT) were conducted to evaluate glucose tolerance and insulin sensitivity, as previously described (20). At the end of the experiments, various tissues, including all fat pads, were dissected, weighed, and harvested for further analyses, including mRNA expression, protein expression, and immunohistochemistry.

### Quantitative reverse transcriptase (RT)-PCR

Quantitative RT-PCR analysis of mRNA expression was conducted as we previously described (21,22). Briefly, total RNA was extracted from fat tissue using the Tri Reagent kit (Molecular Research Center, Cincinnati, OH). Levels of mRNAs for target genes were measured using a one-step quantitative RT-PCR protocol with the TaqMan Universal PCR Master Mix kit (ThermoFisher Scientific, Waltham, MA) on an Applied Biosystems QuantStudio 3 real-time PCR system (ThermoFisher Scientific). TaqMan primers and probes for all target genes were purchased from Applied Biosystems (ThermoFisher Scientific).

### Histological analysis

Fat tissues were fixed in 10% neutral formalin, embedded in paraffin, and sectioned into 5 µm-thick slices. These sections were processed for hematoxylin and eosin (H&E) staining as we previously described (21).

### Statistics

Data were presented as Mean ± SEM. Different groups in each experiment were compared for difference by one-way ANOVA or Student’s-t test as appropriate. Statistical significance is accepted at *p <* 0.05.

## Results

### Dnmt3b deficiency in adipocyte progenitor cells ameliorates diet-induced obesity in female mice

In this study, we generated PD3bKO by crossing *Dnmt3b*-floxed mice with Pdgfrα-Cre mice. PDGFRα is a key marker of adipocyte progenitor cells that regulates adipogenesis *in vivo* (23,24). The Pdgfrα-Cre line has been widely utilized for adipocyte lineage tracing and progenitor cell adipogenesis studies (23–27). Deletion of *Dnmt3b* in adipocyte progenitor cells led to a 70% reduction in Dnmt3b mRNA levels in both white adipose tissue (WAT) and brown adipose tissue (BAT) (**Suppl. Fig. 1**). To investigate the role of *Dnmt3b* in diet-induced obesity, we fed PD3bKO mice with a HFD and characterized their metabolic phenotype. Female PD3bKO mice displayed significantly lower body weight compared to their fl/fl littermate controls (**Fig 1A**). Analysis of body composition using a Minispec NMR body composition analyzer revealed decreased body fat mass and increased lean mass in female PD3bKO mice (**Fig 1B**), which was associated with reduced weights of interscapular BAT (iBAT), gonadal WAT (gWAT), and liver (**Fig 1C**). Consistent with decreased fat mass, circulating leptin levels were also reduced in PD3bKO mice (**Fig 1D**).

**Figure 1.**
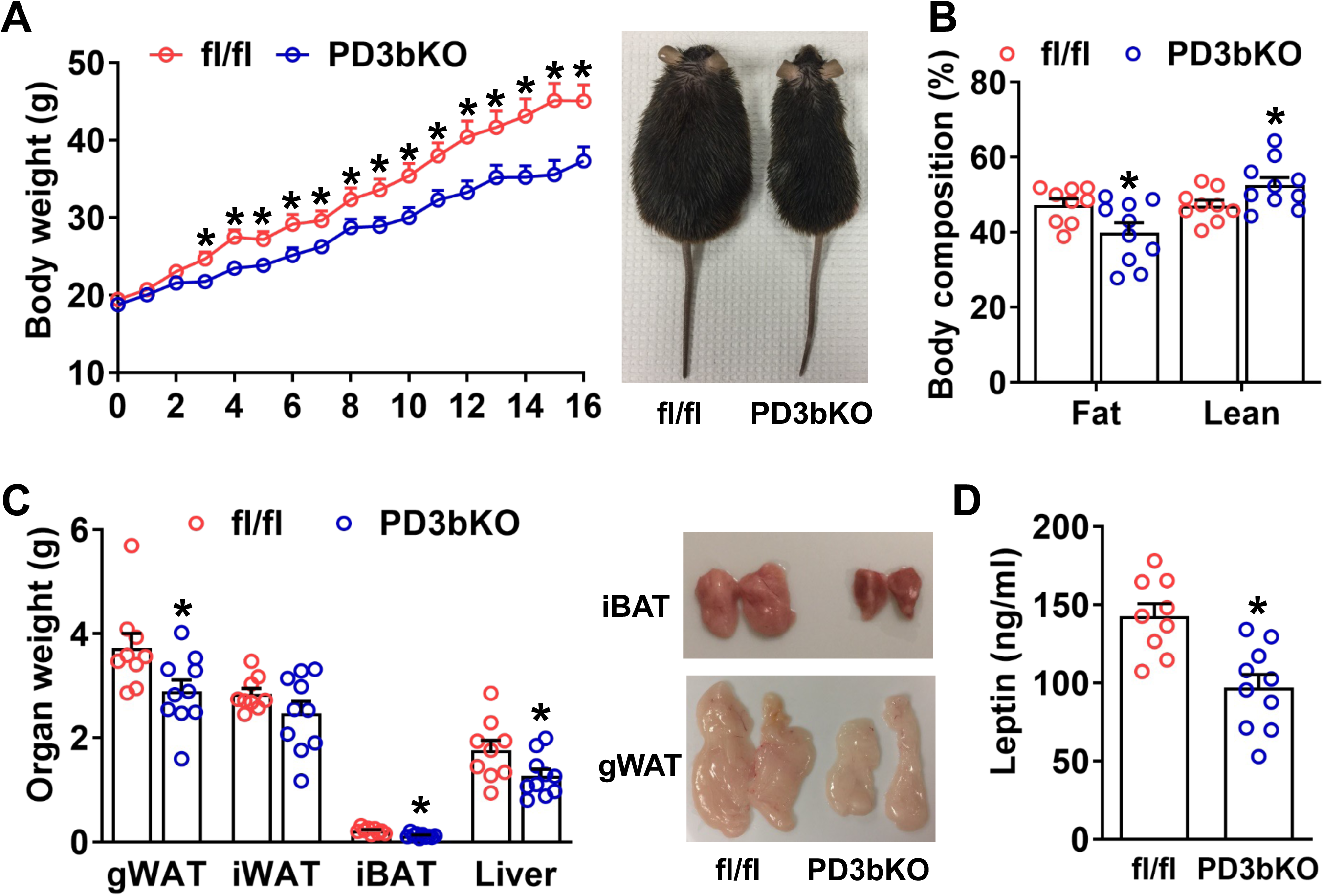
Dnmt3b deficiency in adipocyte progenitor cells prevents HFD-induced obesity in female mice. Six-week-old female PD3bKO mice and their fl/fl littermates were put on an HFD for 16 weeks. (**A**) Body weight growth curve in female PD3bKO and fl/fl mice. (**B**) Body composition measured by a Bruker NMR body composition analyzer in female PD3bKO and fl/fl mice. (**C**) Weights of gonadal WAT (gWAT), inguinal subcutaneous white adipose tissue (iWAT), interscapular brown adipose tissue (iBAT), and liver in female PD3bKO and fl/fl mice. (**D**) Circulating levels of leptin in female PD3bKO and fl/fl mice. All data are expressed as Mean ± SEM; n=9-10/group; \**p* < 0.05 vs. fl/fl.

### Dnmt3b deficiency in adipocyte progenitor cells increases energy expenditure and reduces energy intake in female mice

To identify the factors contributing to the reduced adiposity in PD3bKO mice, we assessed energy expenditure using the PhenoMaster metabolic cage system. PD3bKO mice exhibited higher oxygen consumption (**Fig 2A**), indicating increased energy expenditure. Moreover, a lower respiratory exchange ratio (RER) was observed (**Fig 2B**), suggesting a preference for fat as the primary energy source in knockout mice. Interestingly, we also found reduced food intake in PD3bKO mice compared to fl/fl controls (**Suppl. Fig. 2**). Further analysis of thermogenic gene expression in iBAT using quantitative RT-PCR revealed significant upregulation of thermogenic markers, including *Ucp1, Dio2*, *Pgc1α, Elovl3*, and *Otop1* in female PD3bKO mice (**Fig. 3A**). Consistent with the increased expression of thermogenic genes, H&E staining of iBAT revealed less lipid droplet accumulation in PD3bKO mice than fl/fl littermate controls (**Fig. 3B**). Collectively, these findings suggest that the lean phenotype observed in female PD3bKO mice results from increased energy expenditure and decreased food intake.

**Figure 2.**
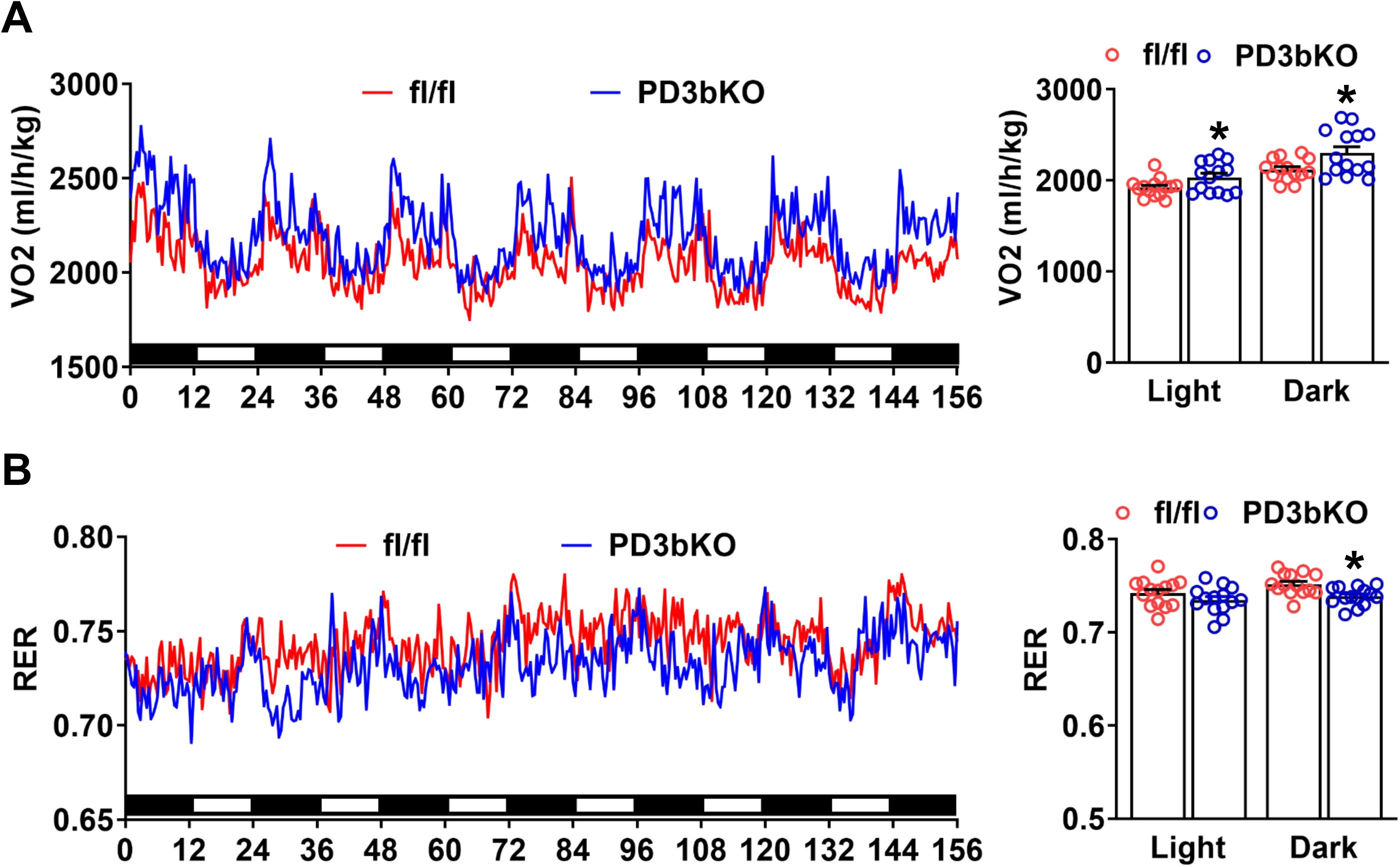
Dnmt3b deficiency in adipocyte progenitor cells promotes energy expenditure in female mice. Female PD3bKO and fl/fl mice fed an HFD were put in TSE PhenoMaster metabolic cage system for metabolic characterization. (**A**) Oxygen consumption (VO2). (**B**) Respiratory exchange ratio (RER). All data are expressed as Mean ± SEM; n=9-10/group; \**p* < 0.05 vs. fl/fl.

**Figure 3.**
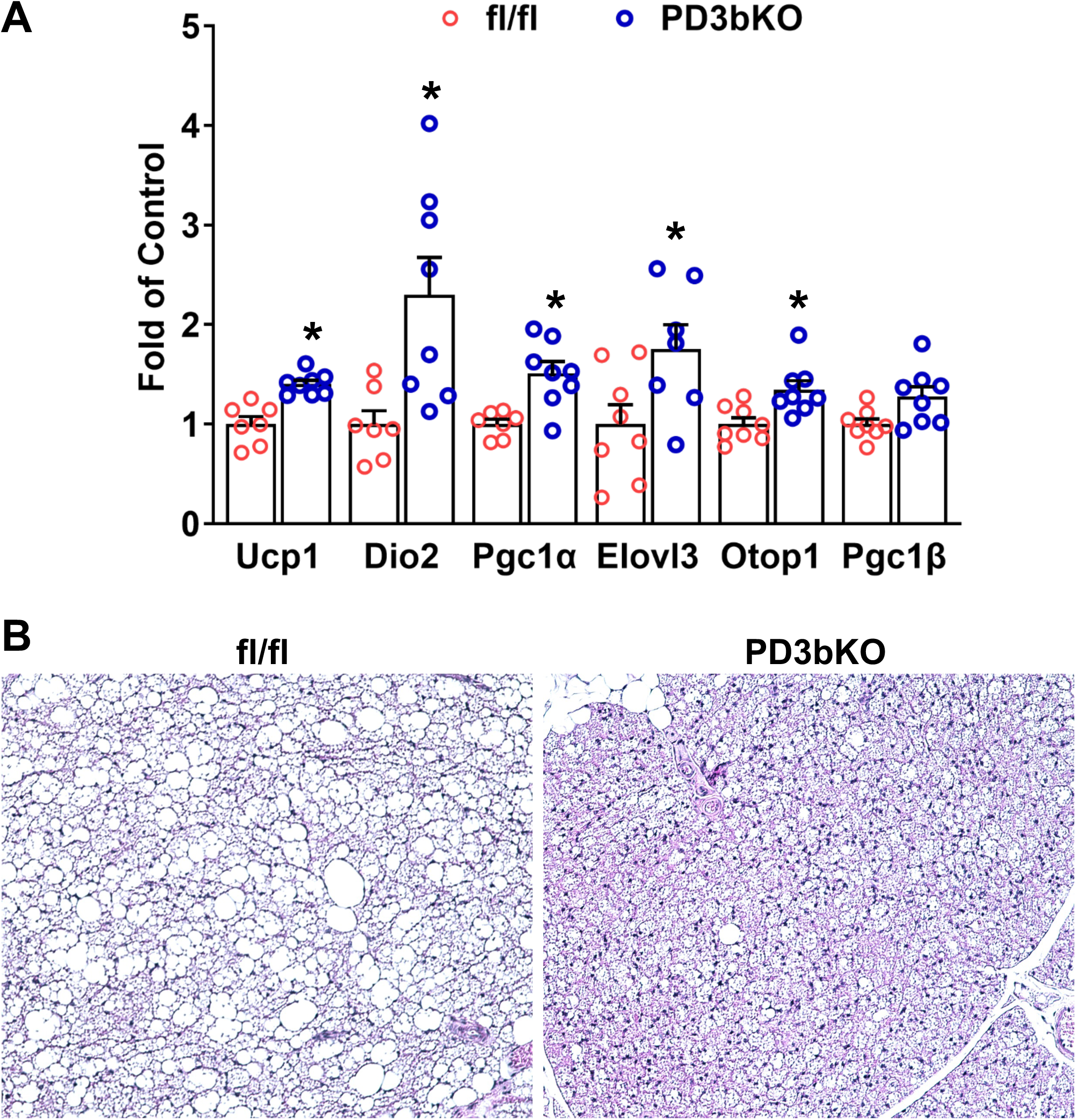
Dnmt3b deficiency in adipocyte progenitor cells promotes brown fat thermogenic program in female mice fed an HFD. Six-week-old female PD3bKO mice and their fl/fl littermates were put on an HFD for 16 weeks. (**A**) Quantitative RT-PCR analysis of thermogenic gene expression in iBAT (n=7-8/group). (**B**) Representative H&E staining images of iBAT. All data are expressed as Mean ± SEM; \**p* < 0.05 vs. fl/fl.

### Dnmt3b deficiency in adipocyte progenitor cells protects against diet-induced gucose intolerance and insulin resistance in female mice

Given the close relationship between adiposity, glucose homeostasis, and insulin sensitivity, we next examined glucose metabolism in PD3bKO mice on HFD. Female PD3bKO mice exhibited lower fasting insulin levels, indicative of improved insulin sensitivity (**Fig 4A**). Consistent with these findings, glucose and insulin tolerance tests revealed enhanced glucose tolerance and insulin sensitivity in PD3bKO mice compared to controls (**Fig 4B** and **4C**).

**Figure 4.**
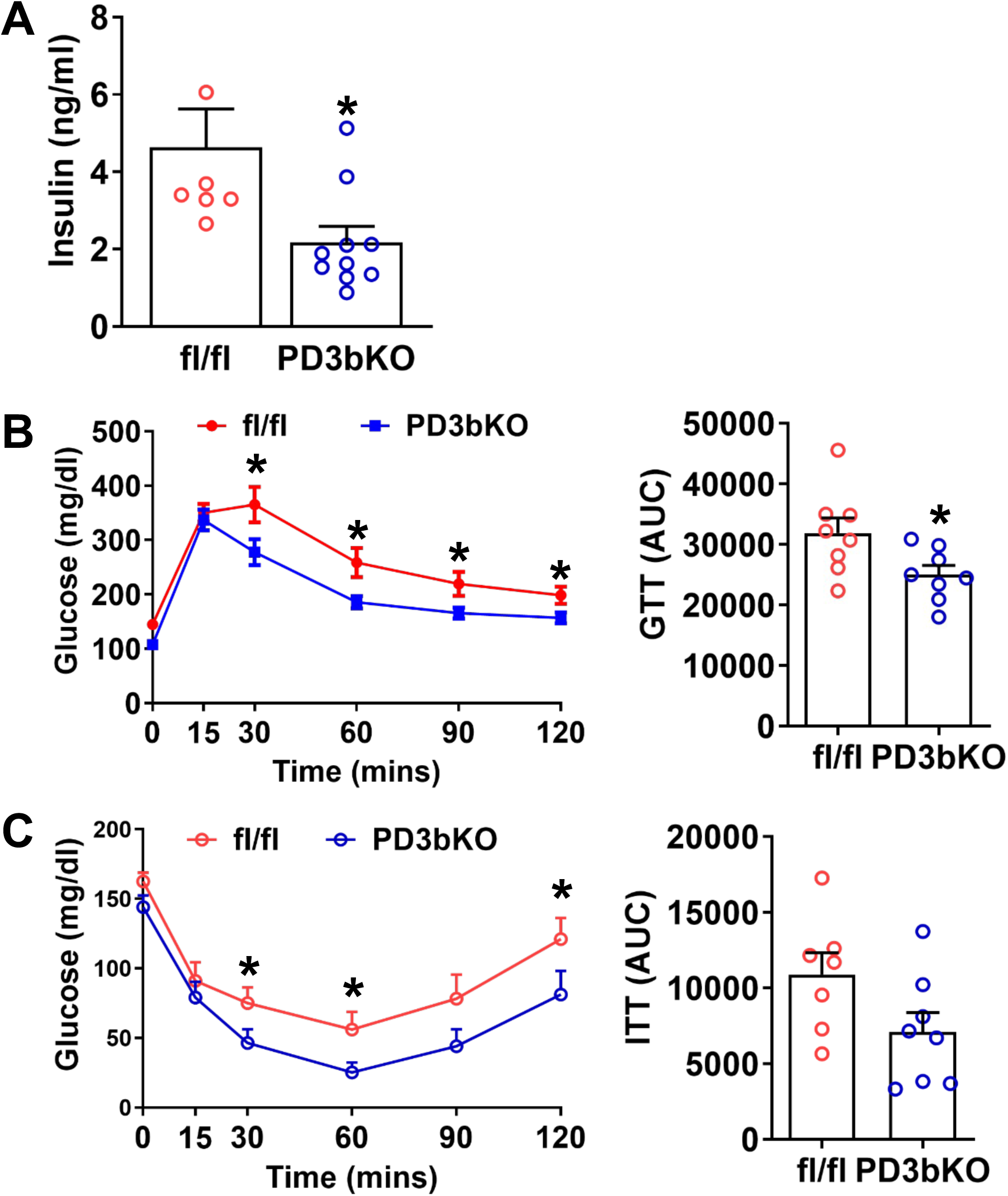
Dnmt3b deficiency in adipocyte progenitor cells improves insulin sensitivity in female mice fed an HFD. Six-week-old female PD3bKO and fl/fl mice were put on an HFD for 16 weeks. (**A**) Circulating insulin levels. (**B**) Glucose tolerance test (GTT) in female PD3bKO and fl/fl mice. (**C**) Insulin tolerance test (ITT) in female PD3bKO and fl/fl mice. AUC, area under curve. All data are expressed as Mean ± SEM; n=8/group; *p<0.05 vs. fl/fl.

### Dnmt3b deficiency in adipocyte progenitor cells impairs insulin sensitivity in male mice

In contrast with the lean phenotype observed in female PD3bKO mice, male PD3bKO mice fed an HFD exhibited a trend toward higher body weight, though it did not reach statistical significance (**Suppl. Fig. 3A**). Moreover, male PD3bKO mice showed no significant changes in body composition or fat pad weight compared to controls (**Suppl. Fig. 3B** and **3C**). While glucose tolerance, assessed by GTT, remained unchanged in male PD3bKO mice (**Suppl. Fig. 3D**), they displayed increased insulin resistance, as evidenced by ITT results (**Suppl. Fig. 3E**). These findings indicate that there is a sexually dimorphic metabolic response to *Dnmt3b* deletion in adipocyte progenitor cells.

## Discussion

In this study, we demonstrated that female PD3bKO mice with *Dnmt3b* deficiency in adipocyte progenitor cells exhibit resistance to HFD-induced obesity and insulin resistance. This phenotype is associated with increased energy expenditure and reduced caloric intake. These findings align with and build upon our prior observations highlighting the significance of epigenetic regulation in obesity and metabolic disease development. The plausibility of this study was derived by prior observations. Emerging evidence has demonstrated epigenetic regulation as a key mechanism mediating the development of obesity and metabolic diseases. Obesity, a multifactorial disease, results from complex interactions between genetic and environmental factors. Environmental influences, such as diet, modulate gene expression through epigenetic reprogramming, thereby serving as a mechanistic link between external factors and disease outcomes (28–32). DNA methylation, a common epigenetic modification, has been implicated in regulating genes involved in various metabolic pathways, including *Ucp1* (33), *Pgc1α* (34,35), *Pparγ* (36), *Lpl* and *aP2* (36), *leptin* (28,37), etc. Our previous work has further explored the role of DNA methylation in metabolic regulation. For instance, we demonstrated that DNA methyltransferases DNMT1 and DNMT3A exhibit stage-specific regulatory effects on 3T3-L1 cell adipogenesis, promoting early differentiation while inhibiting late-stage lipogenesis (14,15). In addition, we also discovered that obesity-induced factors induce DNA hypermethylation at the PPARγ1 promoter via DNMT1, promoting macrophage polarization, inflammation, and the progression of insulin resistance and atherosclerosis (20,38,39). Additionally, we reported that DNMT1 deficiency in neurons protects against diet-induced obesity and insulin resistance (40). Our recent investigations into histone modifications have revealed their role in regulating the thermogenic program of BAT (41–43). These findings motivated us to extend our focus to DNA methylation in BAT development and thermogenic function (16–18). Notably, we found that Dnmt1 or Dnmt3a deficiency in BAT promotes its remodeling into a skeletal myocyte-like phenotype, resulting in decreased energy expenditure and increased adiposity (16,18).

In contrast to these findings, the present study reveals that female PD3bKO mice with *Dnmt3b* deletion in adipocyte progenitor cells exhibit resistance to HFD-induced obesity. Although the precise mechanisms underlying this lean phenotype remain unclear, these mice show increased thermogenic activity in interscapular BAT (iBAT). While it remains uncertain whether PDGFRα is expressed in brown adipocyte progenitor cells, which share a developmental lineage with skeletal muscle, PDGFRα has been identified in UCP1-positive beige progenitor cells (44). PDGFRα-Cre might also be expressed in BAT progenitor cells, potentially leading to *Dnmt3b* deletion in BAT. Consistent with this hypothesis, PD3bKO mice exhibit reduced *Dnmt3b* levels in iBAT, as shown in Supplemental Figure 1. We speculate that early Dnmt3b deletion in brown adipocyte progenitor cells enhances brown fat development and thermogenic function in mature brown adipocytes, thereby increasing energy expenditure and reducing adiposity. This is consistent with our recent report that deletion of *Dnmt3b* in mature brown adipocytes using UCP-1-Cre driver ameliorates obesity in female mice (17). These findings suggest that Dnmt3b may exert distinct roles in brown fat thermogenic regulation compared to Dnmt1 and Dnmt3a (16). Like many epigenetic regulators, DNMTs likely perform stage-specific functions during development. Further studies are warranted to elucidate the precise mechanism by which Dnmt3b regulates adipogenesis in brown adipocyte progenitors.

Additional questions emerge from our observations of female PD3bKO mice. These animals exhibit reduced food intake while maintaining physical activity, both of which contribute to their lean phenotype. It remains to be determined whether these effects stem from peripheral factors, such as fat-derived secretory molecules, or whether they involve potential *Dnmt3b* deletion in neuronal progenitor cells due to PDGFRα-Cre expression in the central nervous system. Future investigations should address these possibilities to further clarify the underlying mechanisms driving the observed phenotype.

Unlike female PD3bKO mice, male PD3bKO mice displayed a trend toward higher body weight and greater insulin resistance when fed an HFD, suggesting a phenotype that contrasts with the protective effects observed in females. This difference may reflect sexual dimorphism in metabolic phenotypes, a phenomenon documented in both humans and rodents. For example, women generally exhibit a higher fat composition compared to men (45) and show distinct patterns of fat distribution, favoring subcutaneous fat storage, while men are more prone to visceral fat accumulation (46). Sexual dimorphism also extends to lipid metabolism, including variations in lipolysis and triglyceride secretion/clearance (47). A well-recognized contributor to these differences is the sex hormone estrogen and its receptors that are known to play a pivotal role in regulating metabolic processes (48). It is plausible that estrogen and its downstream signaling pathways provide additional metabolic protection in female PD3bKO mice on an HFD. Supporting this hypothesis, our previous work identified estrogen receptor demethylation as a potential regulatory mechanism (40). These findings underscore the importance of considering sex-specific factors in metabolic research and highlight the need for further studies to elucidate the role of estrogen signaling in mediating the observed phenotypic differences between male and female PD3bKO mice.

In summary, we generated PD3bKO mice with Dnmt3b deficiency in adipocyte progenitor cells and demonstrated that these mice are resistant to HFD-induced obesity and insulin resistance. This resistance is accompanied by enhanced energy expenditure, increased locomotor activity, and reduced caloric intake, which together likely contribute to the observed lean phenotype. Additionally, Dnmt3b deficiency promotes thermogenic programming in brown fat. Interestingly, male PD3bKO mice exhibit no significant changes in body weight but display reduced insulin sensitivity, highlighting a sexually dimorphic metabolic phenotype in this knockout model. Overall, our findings demonstrate that Dnmt3b in adipocyte progenitor cells plays a crucial role in regulating energy metabolism and body weight, particularly in female mice.

## Acknowledgments

This work was supported by NIH grants R01DK107544, R01DK118106, and R01DK125081 and the American Diabetes Association (ADA) grant 1-18-IBS-260 to BX and by NIH grants R01DK115740 and R01DK118106 and ADA grant 1-18-IBS-348 to HS.

