## Supplemental Figures, Figure Legends and Tables for "Dnmt3b deficiency in adipocyte progenitor cells ameliorates obesity in female mice"

**Supplemental figure legends**

**Supplemental figure 1.** *Dnmt3b* mRNA levels in inguinal WAT (iWAT) and interscapular BAT (iBAT) of female PD3bKO and fl/fl mice (n=7/group). All data are expressed as Mean ± SEM; *$p < 0.05$ vs. fl/fl.

**Supplemental figure 2.** Food intake of female PD3bKO and fl/fl mice fed an HFD. All data are expressed as Mean ± SEM; n=9-10/group; *$p < 0.05$ vs. fl/fl.

**Supplemental figure 3.** Dnmt3b deficiency in adipocyte progenitor cells does not change body weight in male mice fed an HFD. Six-week-old male PD3bKO mice and their fl/fl littermates were put on an HFD for 16 weeks. (**A**) Body weight growth curve in male PD3bKO and fl/fl mice. (**B**) Body composition measured by a Bruker NMR body composition analyzer in male PD3bKO and fl/fl mice. (**C**) Organ weight of epididymal WAT (eWAT), inguinal subcutaneous white adipose tissue (iWAT), interscapular brown adipose tissue (iBAT), and liver in male PD3bKO and fl/fl mice. (D) Glucose tolerance test (GTT) in male PD3bKO and fl/fl mice. (E) Insulin tolerance test (ITT) in male PD3bKO and fl/fl mice. All data are expressed as Mean ± SEM; n=10/group; *$p < 0.05$ vs. fl/fl.

**Supplemental figure 1**

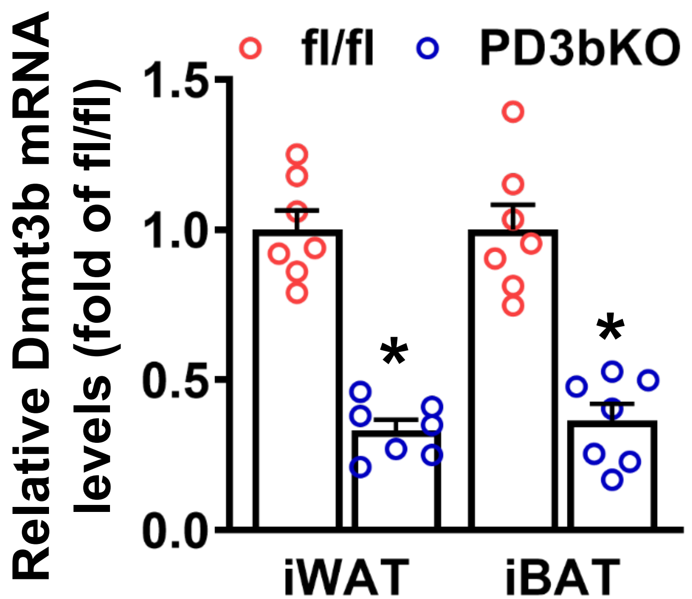

**Supplemental figure 2**

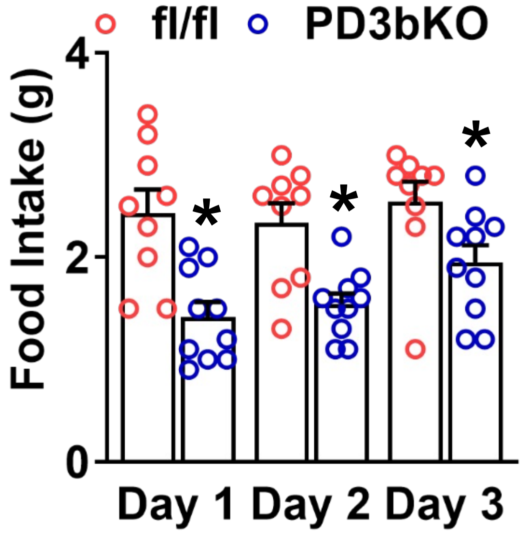

### Supplemental figure 3

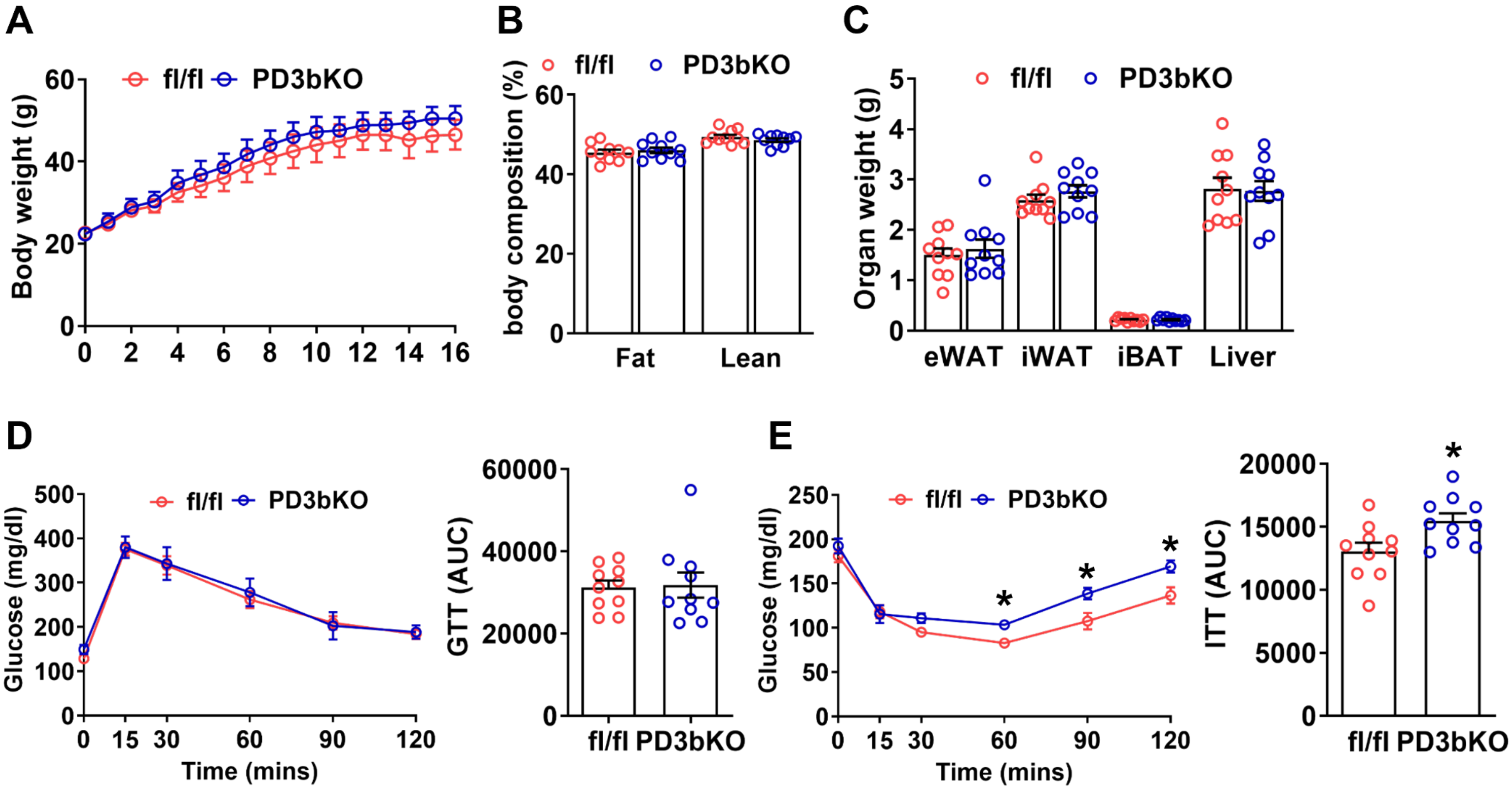
